# RNA polymerase active centre compensates for the absence of transcription proofreading factors in Cyanobacteria

**DOI:** 10.1101/680546

**Authors:** Amber Riaz-Bradley, Katherine James, Yulia Yuzenkova

## Abstract

The vast majority of organisms possess transcription elongation factors, the functionally similar bacterial Gre and eukaryotic TFIIS/TFS. Their main cellular functions are to proofread errors of transcription and to restart elongation via stimulation of RNA hydrolysis by the active centre of RNA polymerase (RNAP). Very few taxons lack these factors, including the large evolutionarily ancient group of cyanobacteria and their descendants, the chloroplasts. How do they cope? What compensatory mechanisms they possess?

We found that cyanobacterial RNAP functionally substitutes for Gre/TFIIS - it does not stall on DNA, it efficiently catalyses the proofreading reaction of RNA hydrolysis, and the drop in transcription fidelity is only fractional, as confirmed by NGS. This alternative, presumably primordial, route to fidelity and processivity in the absence of Gre/TFIIS factors is based on the active site of RNAP stabilisation in a closed conformation. However, here lies a trade off - a severely reduced ability of this active site to recognise regulatory pausing signals. We suggest that perhaps the main advantage of Gre/TFIIS acquisition was to allow transcription regulation via pausing; with increase in fidelity as a bonus side effect.

## Introduction

Correct and fast copying of genomic sequence from DNA into RNA during transcription is vital for faithful expression of genetic information. The elongation stage contributes significantly into the overall efficiency of transcription. Misincorporated and inactivated RNAP molecules are reactivated by special elongation factors via stimulation of transcript hydrolysis. The mechanism of hydrolysis is remarkably similar among all living organisms (1-3). Escape from transcriptional arrest via transcript cleavage is essential for efficient transcript elongation and cell viability (4). In the vast majority of organisms characterised so far, weak intrinsic proofreading by RNAP is augmented by elongation factors, GreA factors in bacteria and TFIIS/TFS in archaea/eukaryotes (5-9). These proteins are not homologous between the two kingdoms of organisms but share a general mechanism. This similarity suggests a functional convergence of proofreading factors and a strong incentive to encode them.

And yet, Gre/TFIIS factors are notably absent from one of the largest groups of bacteria, Cyanobacteria (and several other smaller free-living taxons). These Gram-negative photosynthetic bacteria and evolutionary cousins of chloroplasts, are one of the oldest, most successful and widespread phylogenetic groups. Assuming the represent an evolutionary primitive mechanism of transcription, we can get a glimpse of an alternative way of supporting the fidelity and processivity of transcription, and perhaps the evolutionary reason for the acquisition of additional elongation factors in all other lineages.

In all multisubunit RNAPs transcript elongation and transcript hydrolysis are performed by the same, highly conserved active centre of RNAP. For all reactions, RNAP utilises a two Me^2+^ ion mechanism (10). The first Me^2+^ is held by an invariant triad of aspartate residues, and the second Me^2+^ is brought into the active centre by the substrate: nucleoside triphosphate (NTP), pyrophosphate or hydroxyl ion. Additionally, efficient catalysis of either phosphodiester bond formation or hydrolysis requires the correct folding of a flexible domain of the active centre, the Trigger Loop (TL), and its supporting domain, the Bridge Helix (BH) (6-9). The TL oscillates between closed (active) and open (inactive) conformations via intermediate conformations; most reactions do not require full TL opening (11). During RNA synthesis, closing of the TL stabilises transition state of reaction, providing an induced fit mechanism of catalysis (12-14).

After RNAP incorporates an incorrect NMP by mistake, the 3’-end piece of RNA loses contact with the template, RNAP backtracks one base pair along the template, and the elongation complex becomes deactivated. Elongation resumes once the error-containing piece of RNA is cleaved out, and a new, correctly paired RNA 3’-end is generated. The hydrolysis reaction following misincorporation is very efficient, due to the stabilisation of elongation complexes in a 1bp backtracked conformation (11). This general mode of transcriptional proofreading via transcript hydrolysis is similar among all characterised RNAPs (1-3). The TL participates in the hydrolysis reaction either by positioning of the reactants and stabilising the transition state (12), or directly in some cases (15). The fascinating feature of intrinsic proofreading is the direct involvement of the 3’-end of a transcript in its own excision, resembling ribozymes (16). The involvement and the nature of a general base in catalysis is still a matter of continuing debate in the field (15, 17).

It is generally accepted that intrinsic hydrolysis is not fast enough to correct transcriptional mistakes in real time in the cellular context, and that the modern proofreading mechanism relies on specialised protein factors. The best-studied of these prokaryotic protagonists is *E. coli* GreA. GreA is a member of a group of homologous proteins which bind the vicinity of the secondary channel. This channel provides a route for substrate entry into the active centre (hence their alternate name, secondary channel binding factors). By inserting a coiled-coiled domain through the channel into the active centre, GreA flips the TL open and physically replaces it in the active site, thereby stopping elongation. Acidic amino acid residues at the tip of the coiled-coil domain of GreA stabilise the second Mg^2+^ ion and coordinate a water molecule greatly increasing the efficiency of hydrolysis (18-20). As a result GreA improves the fidelity of transcription by up to two orders of magnitude in some instances (21). Gre factors can also reactivate correctly paired complexes that have backtracked and arrested for different reasons (such as prolonged pauses or DNA lesions) (22, 23). Consequently, timely cleavage and reactivation of backtracked paused transcription complexes is a vital mechanism to remove stalled RNAPs out of the way of the replisome to avoid collisions, and to prevent the formation of traffic jams of RNAPs on actively transcribed genes in bacteria (24, 25). Secondary channel binding factors are not essential in laboratory conditions, but their loss is severely detrimental for viability of all bacterial species tested, including *E.coli* and *Streptococcus pneumoniae*; and their importance rises in different stress conditions (24, 26).

Here, we studied the native RNAP of a widely used research and biotechnology species of cyanobacterium, *Synechococcus sp* PCC 6803 (*Ssp*RNAP). Most of our results were also reproduced with the RNAP of the distantly related *Synechococcus elongatus* 7942, (*Sel*RNAP), another model cyanobacterium, which allowed a generalisation of our findings for the whole group. We found that cyanobacterial transcriptional fidelity is not severely compromised by the absence of proofreading factors, and that the level of *in vivo* mistakes in mature RNA is only fractionally higher than that of *E. coli. In vitro Ssp*RNAP is not more accurate in substrate choice, yet proofreads transcription up to two orders of magnitude faster than *Ec*RNAP. We suggest that the *Ssp*RNAP active site tends to reside in a closed, hydrolytically-competent conformation. In cyanobacteria, the hydrolysis reaction is assisted by a general base, similarly to the Gre-stimulated reaction, and in contrast to *Ec*RNAP. Another consequence of the cyanobacterial active site conformation is the suppression of transcriptional pausing and termination.

## Results

### Absence of Gre results in a moderate drop in the fidelity of transcription

Since Gre factors are a major contributor to fidelity, does an absence of Gre factors result in a higher level of mistakes in RNA? To answer this question we assessed the levels of mismatches in the mature RNA *in vivo* by Next Generation Sequencing of the transcriptomes of *E. coli* and *Synechocystis sp* 6803. To minimise the level of technical mistakes, we used the reverse transcriptase with highest fidelity available, which had been employed previously to determine the *in vivo* rates of transcriptional mistakes in *E. coli* (27). We compared the rate of mismatches at a particular position in the middle of the sequencing reads and found that levels of some base changes are indeed higher in *Synechocystis* (C to U, and A to G (Fig. 1)), but at most increased to 125 percent. This increased level of *in vivo* mistakes in RNA although does not remove the probability entirely, argues against any additional unknown proofreading factors in *Synechocystis sp* 6803.

**Figure 1.**
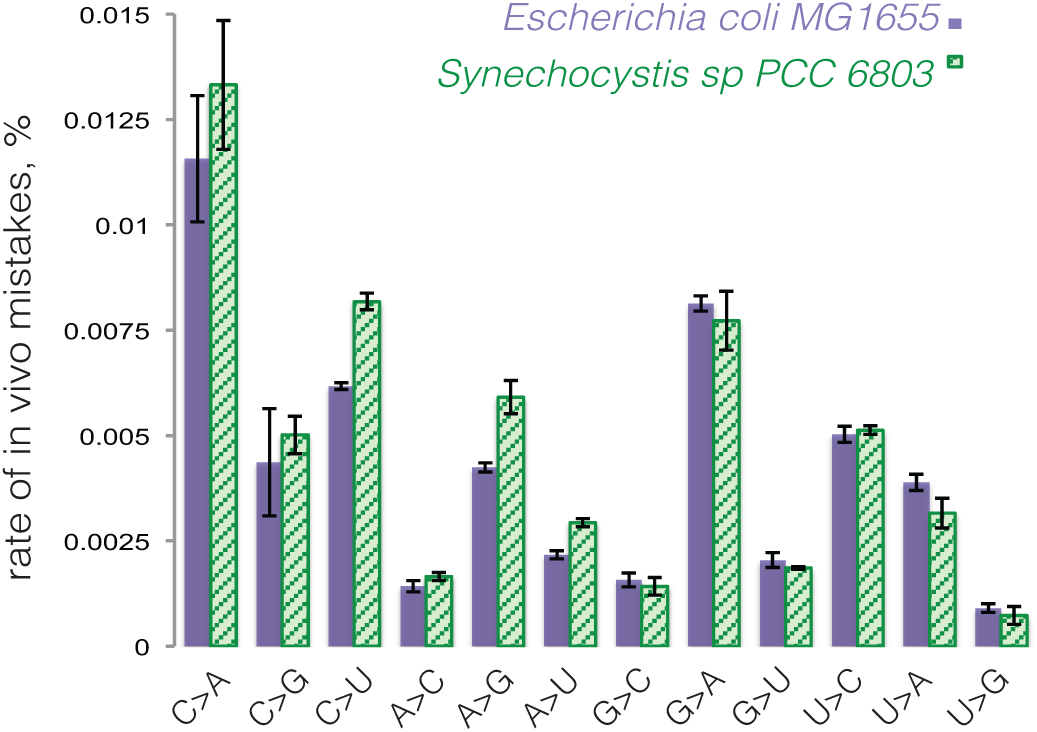
Rates of *in vivo* transcriptional mistakes are comparable between *E.coli* MG1655 and *Synechocystis sp* PCC 6803. The graph shows percentages of total specific *in vivo* mistakes in mature transcripts for *E. coli* MG1655 and *Synechocystis sp* PCC 6803, in specific position of the sequencing read, calculated based on Next Generation Sequencing of total RNA from two species (See Methods for details). Error bars represent standard deviation from biological triplicates.

### *Ssp*RNAP is not slow or accurate, but has high proofreading activity

We assumed that cyanobacteria compensate for the proofreading factors’ absence by having either more accurate incorporation, or more efficient error proofreading (or possibly both). It has been suggested that cyanobacterial RNAP is a slow elongating enzyme (28), which could contribute to its accuracy by providing a longer time frame for correct substrate selection. We found that this is not the case, judging from single nucleotide addition experiments in *in vitro* assembled elongation complexes, ECs. As can be seen from plot on Fig. 2A, the rates of substrate addition were comparable for *Ssp*RNAP and *Ec*RNAP in four elongation complexes on template 1.

**Figure 2.**
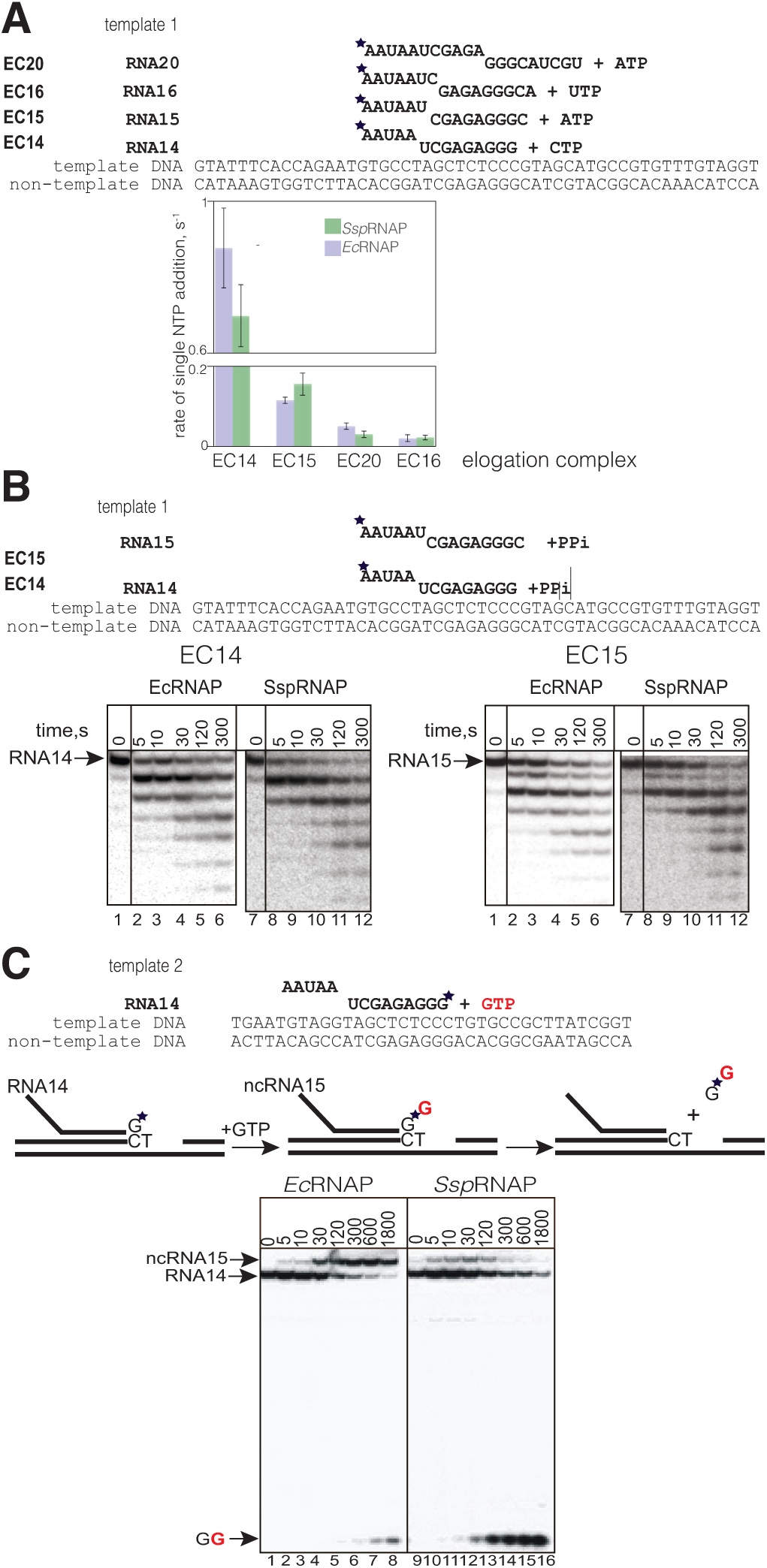
A. Rates of incorporation of correct NTPs are similar for SspRNAP and EcRNAP. Rates of incorporation (s-1) of the single correct (1μM) substrates by SspRNAP and EcRNAP in elongation complex with 14nt, 15nt and 16nt and 20nt long RNA (EC14, EC15, EC16 and EC20). These complexes were obtained by extension of initial RNA13, labelled with 32P at 5’ end, upon NTPs addition. Error bars represent standard deviations from triplicate experiments. Kinetics of pyrophosphorolysis in elongation complexes EC14 and EC15 by EcRNAP and SspRNAP. **B. Rates of pyrophosphorolysis are similar for SspRNAP and EcRNAP**. Kinetics of pyrophosphorolysis in EC14 and EC15 on template 1 at 250 μM pyrophosphate. **C. SspRNAP misincorporates substrate with same efficiency as EcRNAP, and proofreads a mistake faster.** Kinetics of misincorporation by EcRNAP and SspRNAP, representative gel for misincorporation reaction of GTP instead of template-dictated ATP in EC14. Schematics above the gel shows elongation complex and reactions of misincrorporation and subsequent hydrolysis with dinucleotide product release. Asterisk indicates that RNA is labelled at the 3’ end, which allows monitoring of misincorporation and proof-reading simultaneously. Initial 14nt RNA (RNA14) after GTP misincorporation elongates to ncRNA15, then 3’ incorrect dinucleotide piece of the transcript (GG) is cleaved out.

The rates of the pyrophosphorolysis reaction, a direct reversal of nucleotide addition, were similar for the two enzymes in EC14 and EC15 (Fig. 2B). In addition to having the same rate of catalysis, NTP addition and pyrophosphorolysis experiments demonstrated that *Ssp*RNAP is at a similar equilibrium point between pre- and post-translocation states to that of *Ec*RNAP, since these reactions can proceed exclusively from a post- or pre-translocated state, correspondingly.

To test how readily *Ssp*RNAP incorporates incorrect substrates, we tested kinetics of misincorporation of noncognate 1mM NTPs, which is within the range of cellular concentration, into 14 nt long 3’-end labelled RNA in assembled elongation complex EC14 on template 2 (Fig. 2B and Fig. S1). This set up allows simultaneous observation of both misincorporation and proofreading via dinucleotide cleavage. In the experiment shown in Fig. 2C RNAP was forced to incorporate GTP instead of template-dictated ATP. The rate of misincorporation, calculated as rate of initial RNA14 transition into reaction products, was slightly higher for the *Ssp*RNAP (0.016 s^−1^ compared with 0.012 s^−1^ for *Ec*RNAP), suggesting that *Ssp*RNAP is not more accurate. Notably, however, the amount of erroneous transcript, *nc*RNA15 for *Ssp*RNAP was significantly lower at all time points, due to very efficient cleavage of the erroneous 3’-end dinucleotide, pGpG (compare the dinucleotide bands in lanes 6-8 with 14-16, Fig. 2C). A similar effect on miscincorporation and cleavage was observed for misincorporation of CMP instead of UMP (Fig. S2).

We conclude that *Ssp*RNAP has highly efficient proofreading activity, rather than highly accurate incorporation. Therefore, it appears that efficient intrinsic hydrolysis is a primary compensatory mechanism for the Gre factors’ absence in cyanobacteria.

### The molecular mechanism of fast *Ssp*RNAP hydrolysis

What is the molecular mechanism behind fast hydrolysis exhibited by *Ssp*RNAP? Efficient hydrolysis requires particular geometry of the reactants – scissile phosphate bond, two Mg^2+^ ions, and attacking water. Since the 3’-end NMP of the RNA provides additional chemical groups to the active center, the characteristics of the reaction also depend on the nature of this NMP (16).

We investigated all elements of the hydrolysis mechanism, using *m*isincorporated *e*longation *c*omplexes, mECs – elongation complexes where the 3’ end NMP of the RNA is non-complementary to the template base while the DNA template and non-template strands were fully complementary to each other. These complexes mimic the state of elongation complexes after misincorporation, which is one of the main targets of Gre factors in the cell. These elongation complexes are stabilised in a 1bp backtracked conformation, which removes any input from backtracking into the rate of second phosphodiester bond hydrolysis (16). We tested elongation complexes of 15 nt RNA with either U at the 3’-end (mispaired with template G) or A (mispaired with template C), mEC(U) and mEC(A) respectively, (schemes of the reaction are above the graphs on Fig. 4A, 4B and sequences on Fig. S1). In these elongation complexes, the *K*_M_ for Mg^2+^ was similar for *Ssp*RNAP and *Ec*RNAPs at pH 7.9 in both cases, but *k*_cat_ values for *Ssp*RNAP were 30 and 53 times higher than *Ec*RNAP for mEC(U) and mEC(A), respectively. These results suggest that the increased rate of hydrolysis does not come from the stabilisation of the second Mg^2+^ ion in the active site (the mechanism proposed for Gre factors), which is consistent with both enzymes having conserved amino acid residues in the vicinity of the catalytic Mg^2+^ ions.

**Figure 3.**
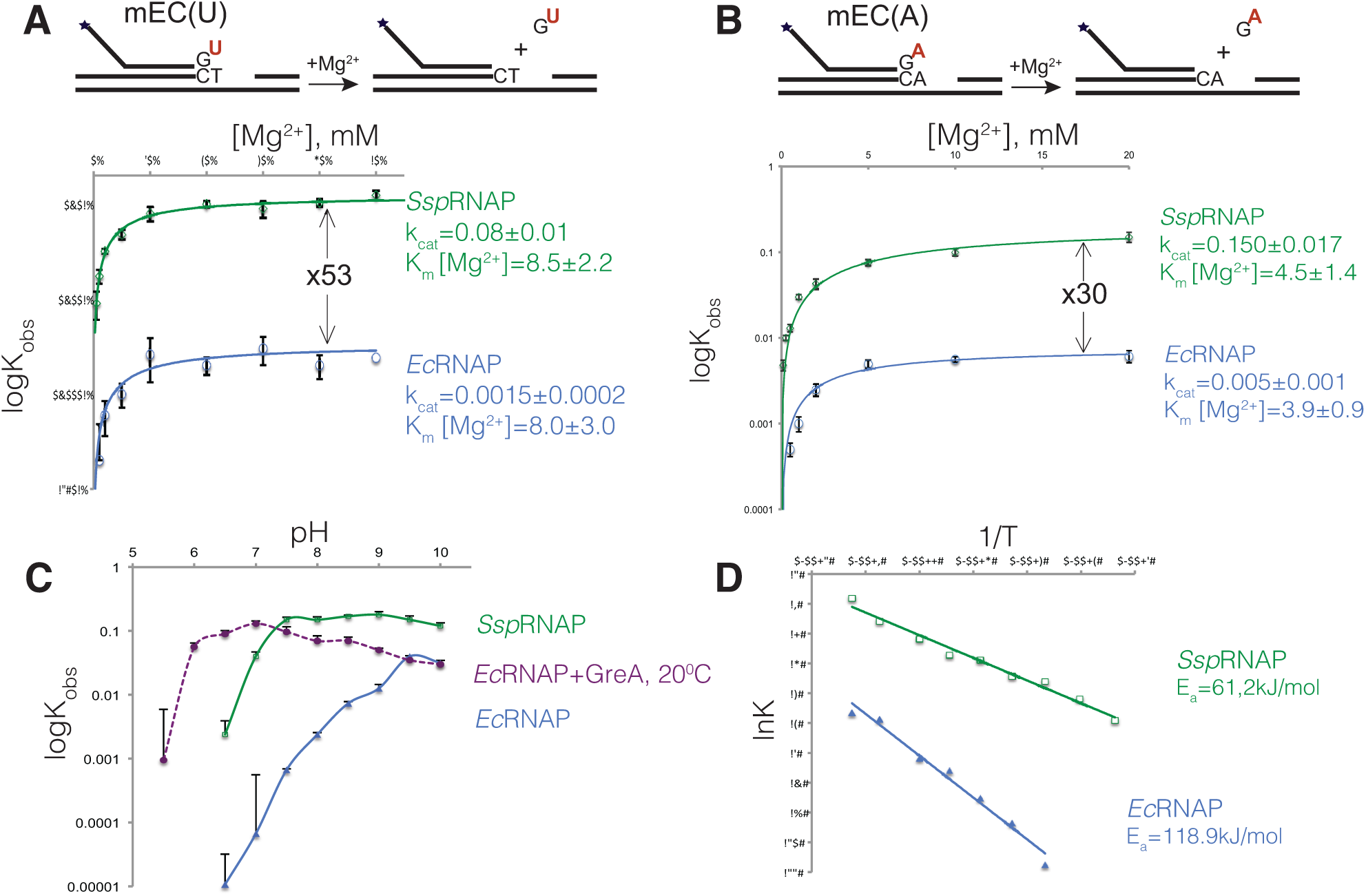
Molecular mechanism of fast transcript hydrolysis by SspRNAP. A and B. Mg^2+^ dependencies of the hydrolysis rate of the second phosphodiester bond in mEC(U) and mEC(A), respectively, by Ec and SspRNAPs. Schematics above the plots show the elongation complex structures and hydrolysis reaction it undergoes; asterisk indicates that RNA is labelled at the 5’ end. Solid lines represent the graphical fits of data (using SigmaPlot software) to the Michaelis-Menten equation. The k_cat_ (reaction rate in saturating Mg^2+^) and K_M_ [Mg^2+^](23) values are shown next to the plots. Note that error bars are not correctly represented due to logarythmic scale. **C. pH profiles of second phosphodiester bond hydrolysis in mEC(A) complex for intrinsic hydrolysis reaction by EcRNAP, GreA assisted hydrolysis by EcRNAP (at 20°C) and SspRNAP.** The data points are averages of 3 independent experiments, negative error bars are omitted, since they are not correctly represented on logarythmic scale (standard deviation for each experimental point were within 10-15% value). **D. Arrhenius plots for EcRNAP and SspRNAP**, graph fit of lnK to 1/T data to linear equation are shown as straight line, apparent activation energy calculated from equation lnK=lnA-Ea/R(1/T) is shown on the plot. The data points are averages of two independent experiments.

**Figure 4.**
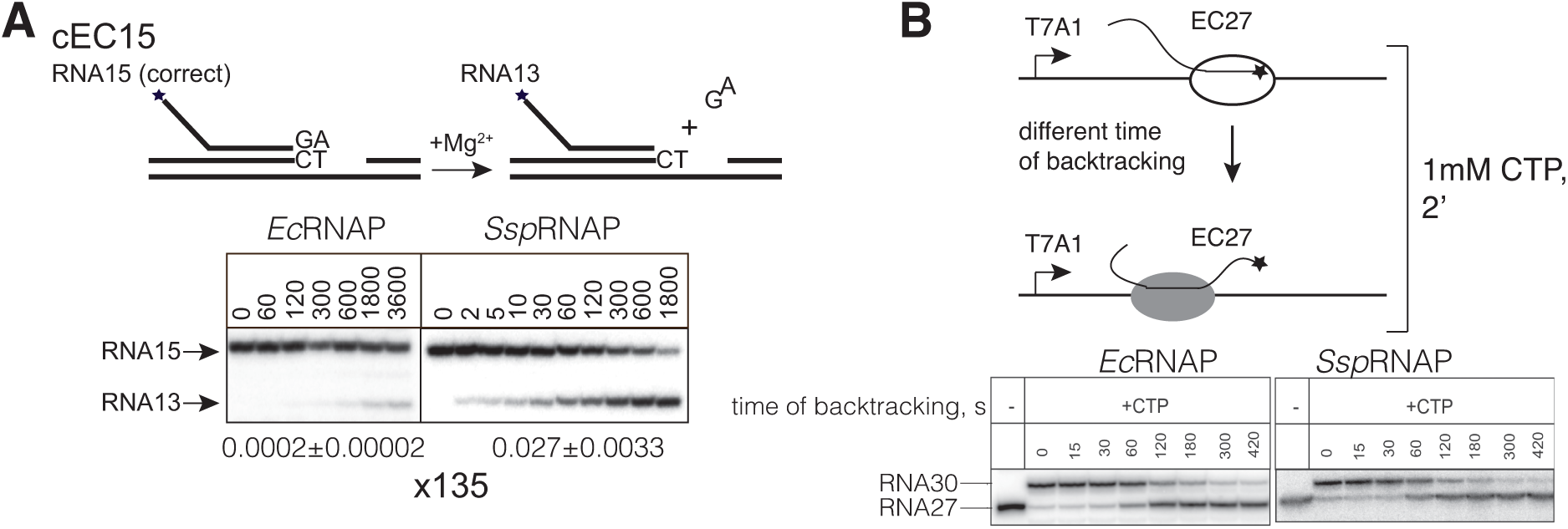
SspRNAP backtracks faster than EcRNAP for 1bp, but not over the long distance. A. Kinetics of hydrolysis of RNA15 in the correct elongation complex (scheme above the gel) by SspRNAP and EcRNAPs with rates of reaction underneath the gel calculated after fitting the data into exponential equation by SigmaPlot software. **B. Backtracking on T7A1 promoter template by SspRNAP and EcRNAP**. Elongation complex EC27 prone to backtracking was formed and incubated at 37°C at pH 6.0 for the indicated periods of time to allow backtracking to happen. Remaining activity was accesses by the ability of complexes able to elongate to position 30 after addition of 1mM CTP.

Hydrolysis requires deprotonation of water, and its efficiency depends on pH of the reaction. The pH-dependence profile of *Ec*RNAP’s rate of intrinsic hydrolysis was drastically different from both *Ssp*RNAP intrinsic hydrolysis and from GreA-assisted hydrolysis by *Ec*RNAP on mEC(A) (Fig. 4C). For *Ec*RNAP in the range of pH 6.5 to 9.7 the dependence is log-linear with a gradient of approximately 0.9, most likely reflecting water ionisation. In contrast, both *Ssp*RNAP and GreA-dependent *Ec*RNAP reactions behave somewhat similarly – the graphs quickly plateau, although at different pH values. We suggest that hydrolysis by *Ec*RNAP is unassisted by protein, in agreement with the work of Mishanina et al., (17). In contrast, in both *Ssp*RNAP and GreA-dependent reaction, a general base activating water is apparently involved,. For the GreA-dependent hydrolysis, the pK_a_ is below 5.5, which most likely corresponds to the pKa of GreA’s active site glutamate (pK_a_ 4.1). For *Ssp*RNAP, the pKa is approximately 6.8, suggesting either a histidine residue (pK_a_ 6.0) or, alternatively, a phosphate group of the transcript (pK_a2_ about 7.2) involvement. There is a possibility of more than one group participation, since the slope of this curve is greater than 1. These results are in line with previous work on *T. aquaticus* RNAP, where a general base was provided in some instances in the form of the Trigger Loop His1242 residue (15).

By analysing temperature dependence of the hydrolysis reaction in mEC(U), we found that the activation energy of the reaction for *Ssp*RNAP is approximately twice lower, compared to *Ec*RNAP (Fig. 4D), suggesting easier isomerisation into a reactive conformation. The same results were observed for mEC(A) (Fig. S5).

### 1bp backtracking, but not long range backtracking, is enhanced in Cyanobacteriam

Misincorporated complexes are not the only targets of Gre factors. The other targets are correctly paired elongation complexes that are left in a backtracked state after arrest or pause, for various reasons (22, 29). For these complexes, the speed of enteri the backtracked state contributes to the overall rate of reaction. Does *Ssp*RNAP backtrack faster by 1bp in a correctly paired elongation complex? We analysed the hydrolysis of a transcripts in a correctly paired elongation complex with 15 nt long RNA with A at the 3’-end, cEC15 (Fig. 5A). Indeed, the overall reaction in the correctly paired transcript in EC15 was faster in *Ssp*RNAP, and the difference in comparison to *Ec*RNAP larger than in misincorporated complexes (Fig. 5A) −135 fold higher in correctly paired elongation complex compared to 30 times in misincorporated mEC(A) (Fig. 3B). We suggest that this higher difference is due to input from faster isomerisation into a 1bp backtracked state by *Ssp*RNAP, but this then rises the question of whether *Ssp*RNAP backtracks faster in general.

**Figure 5.**
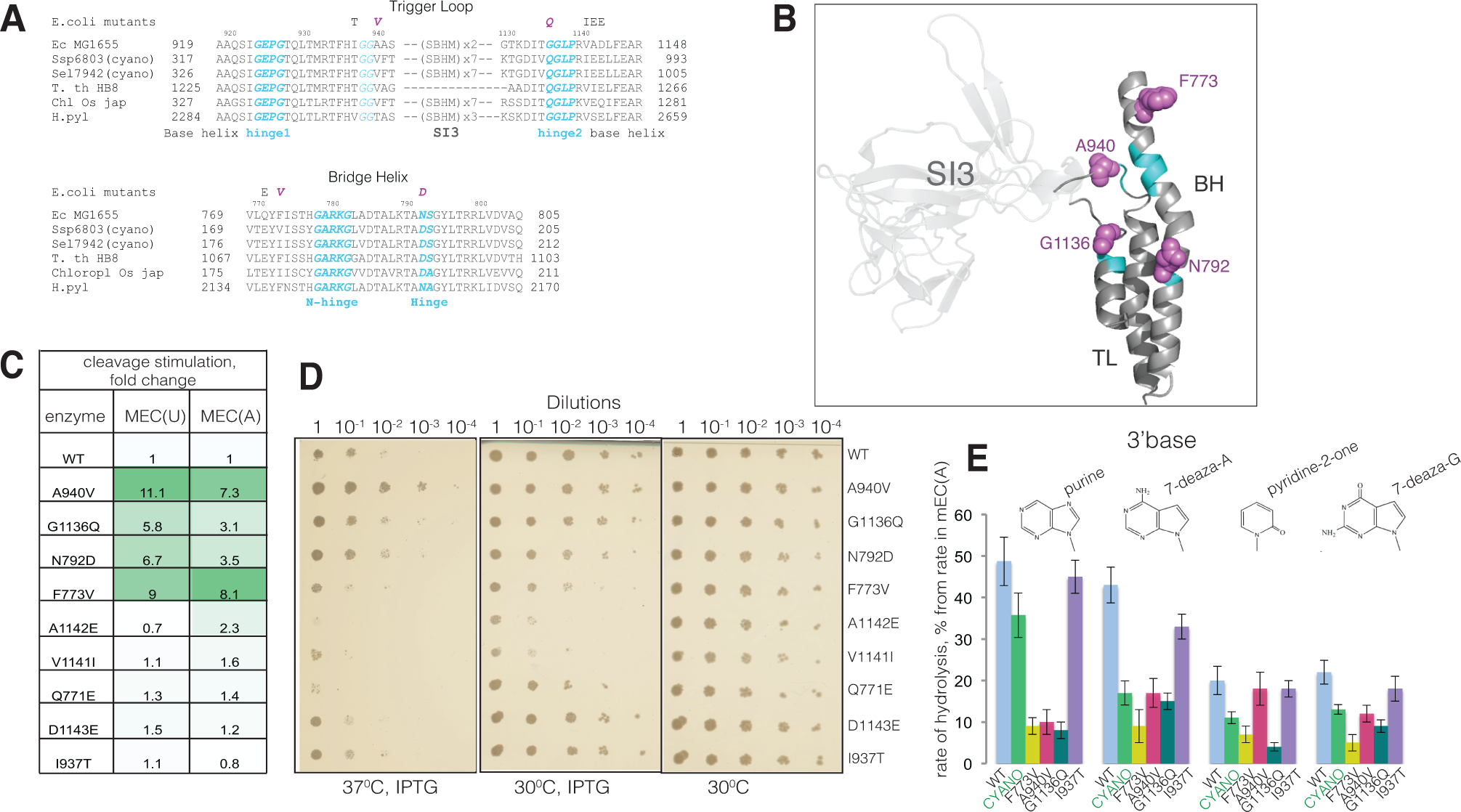
Cyanospecific amino acid residues in TL and BH contribute to the fast transcript hydrolysis by SspRNAP. A. Alignment of the amino acid sequences of the Trigger Loop and part of Bridge Helix of *Synecho-cystis sp* PCC 6803, *Escherichia coli* MG1655, Thermus thermophilus HB8, *Oryza sativa* chloroplast and *Bacillus subtilis*. **B. Structure of *E. coli* RNAP TL and BH domains** from PDB 5IPM aminoacid residues whose substitutions stimulated transcript hydrolysis are shown in magenta. Flexible hinges and additional GG motif in TL are shown in cyan. C. Stimulation of transcript hydrolysis by the changes to cyano - specific amino acid residues in EcRNAP in MEC(U) and MEC(A). Table lists the fold changes of the rates of hydrolysis by the mutant RNAPs in comparison to the WT EcRNAP. D. **Expression of mutant rpoC coding for substitutions A940V, G1136Q, and N792D supresses growth defect of *E. coli* MG1655 ΔgreAgreB strain.** Serial dilutions were plated on Petri dishes and grown over-night at 300C, 300C with IPTG or at 370C with IPTG. **E. Effect of changes of chemical groups of the 3’ RNA base (base structures on top of plot) on hydrolysis rate** by WT EcRNAP (sky blue), SspRNAP (grass green) and mutant EcRNAPs with substitutions F773V (mustard), A940V (hot pink) G1136Q (malachite) and I937T (violet). Plot repre-sent residual activity percentage in elongation complexes with 3’ end RNA modified bases in comparison to canonical base in mEC(A). The error bars represent standard deviation from triplicate data points.

To compare the ability of the RNAPs to backtrack over longer distances we used a well-characterised, prone to backtracking elongation complex with 27 nt long RNA, formed on linear DNA template containing T7A1 promoter. Prolonged incubation of this elongation complex at 37° C typically leads to accumulation of backtracked inactive complexes (30) (Fig. 4B). We monitored transition into a backtracked state by observing a progressive loss of ability of RNAP to extend 27 nt RNA with the addition of NTP substrates. *Ec*RNAP and *Ssp*RNAP were allowed to backtrack for the time intervals indicated in Fig. 3B and supplied with 1mM CTP to elongate RNA in complexes still active from 27 nt to 30 nt long. The fraction of active complexes decreased similarly over time for both enzymes hence we concluded that beyond 1 bp *Ssp*RNAP moves backwards at the same rate as *Ec*RNAP (Fig. 4B). In other words, only a 1bp backtracking, associated with proofreading, is specifically increased in cyanobacteria.

Which parts of the RNAP active site are responsible for efficient cleavage? The fast cleavage observed for *Ssp*RNAP is a hallmark of a closed active site conformation (31). The main flexible domains of the active site are the TL and BH, and both have been previously shown to influence transcript hydrolysis (15, 17, 32). Therefore, it is very probable that conformation and dynamics of these domains are responsible for the efficient transcript hydrolysis characteristic for *Ssp*RNAP. We hypothesised that this efficiency has been achieved by the strategic placement of cyanobacteria-specific amino acid residues in the “hinges” of the TL and/or BH.

### Cyano-specific amino acid residues in the Trigger loop and Bridge Helix stimulate hydrolysis by *Ec*RNAP and suppress the phenotype of Δ*greA*Δ*greB* strain

The trigger loop consists of two helical parts separated by N-terminal and C-terminal “hinges”, and an SI3 insertion, present in both cyanobacteria and *E. coli* (Fig. 5A and 5B). The amino acid sequence corresponding to N-terminal part of the TL is the same for both *Ssp*RNAP and *Ec*RNAP, however several cyanobacteria-specific amino acids can be found in the unstructured region and in the C-terminal base helix of the TL (Fig. 5A and 5B). To investigate if any of these amino acid residues contribute to efficient hydrolysis, we tested mutant *Ec*RNAPs whose native amino acid residues were changed to the cyano-specific ones in the TL (*E. coli* numbering) – I937T, A940V, A941F, K1132G, T1135V, G1136Q, V1141I, A1142E, D1143E and F1145L (Fig. 4A). Rates of RNA hydrolysis were analysed in the assembled misincorporated elongation complexes mEC(A) and mEC(U) (Fig. S1). Most substitutions did not affect the rate of hydrolysis (Fig. 4C and SI Table 1, which also includes additional data on mEC(C)). However, two changes, A940V and G1136Q, increased the rate of hydrolysis by mutant *Ec*RNAP by 7-11 fold and 3-6 fold respectively, in both mECs. The amino acid residues at these positions are too far from 3’-end of RNA to participate in cleavage reaction directly. Notably, A940V is located next to double glycines in the N-terminal part of TL, and the G1136Q substitution changes a flexible glycine into a more rigid glutamine in the C-terminal hinge region of the TL. Both these substitutions have the potential to affect folding dynamics of the TL. Since folding of the TL proceeds in concert with the BH, and changes in the BH affect transcript hydrolysis (32), we looked for additional cyanobacterial-specific amino acid substitutions in the BH. The most conspicuous change is F773V, located close to N-terminal glycine hinge of the BH (Fig. 5A). This substitution is severely detrimental to the growth of *E. coli* (33). F773V *Ec*RNAP increased the rate of the hydrolysis reaction on both mEC(A) and mEC(U) by 8-9 fold (Fig. 5C). Another BH substitution, N792D, increased the hydrolysis rate 3.5-6.5 fold, while Q771E did not have an effect on hydrolysis rate (Fig. 5C).

Could mutant RNAPs with increased proofreading efficiency suppress the temperature sensitive phenotype of an *E. coli* strain with both GreA and GreB factors deleted (23)? To address this question we expressed mutant (F773V, N792D, I937T, A940V, D1143E, Q771E, G1136Q, V1141I, A1142E) or WT β’ subunits from a pRL663 plasmid in the Δ*greA*Δ*greB* MG1655 strain of *E. coli*, plated and grew culture dilutions on solid media at either the permissive (30°C) or the nonpermissive (37°C) temperature with addition of IPTG to induce expression of the mutant subunits. Mutants with increased transcript hydrolysis efficiency (in particular A940V) were able to moderately promote growth in comparison to WT or other, neutral mutants (Fig. 5D). The exception was F773V, which has been previously characterised as generally detrimental for viability of *E. coli* (34).

The 3’-end of the RNA contributes into the efficiency of its own hydrolysis reaction by providing additional coordination groups to water and Mg^2+^ ions (16). Change or removal of these chemical groups of the transcript’s 3’-end base reduces efficiency of hydrolysis. We found that *Ssp*RNAP and hydrolytically proficient, but not hydrolytically neutral, *Ec*RNAPs mutants are more sensitive to chemical modifications of the 3’-base. As can be seen from Fig, 4E, changing 3’-adenine in a mEC(A) to a purine, pyridine-2-one, 7-deaza-A, or 7-deaza-G has a greater effect on the hydrolysis rate (Fig. 4E) of *Ssp*RNAP, and of F773V, A940V and G1136Q *Ec*RNAPs. We suggest that a greater reduction of rate is related to a stronger original mechanism and that a specific folded TL conformation provides some interaction with mismatched 3’-end of RNA, as was proposed by Larson et. al, (35).

### *Ssp*RNAP inefficiently recognises pausing and termination signals

Pausing of RNAP during elongation is accompanied by TL opening (12, 13, 31, 36). We hypothesised that pausing efficiency might be lower for *Ssp*RNAP, due to the tendency of its active site to reside in a closed conformation. In the experiment on Fig. 6A, we performed kinetics of transcript elongation in an assembled elongation complex with 14nt long RNA, EC14, upon addition of a low concentration of all four NTPs. Indeed, the propensity of *Ssp*RNAP to make fewer ubiquitous pauses during elongation, and to reach the end of template faster, is evident from Fig. 6A.

**Figure 6.**
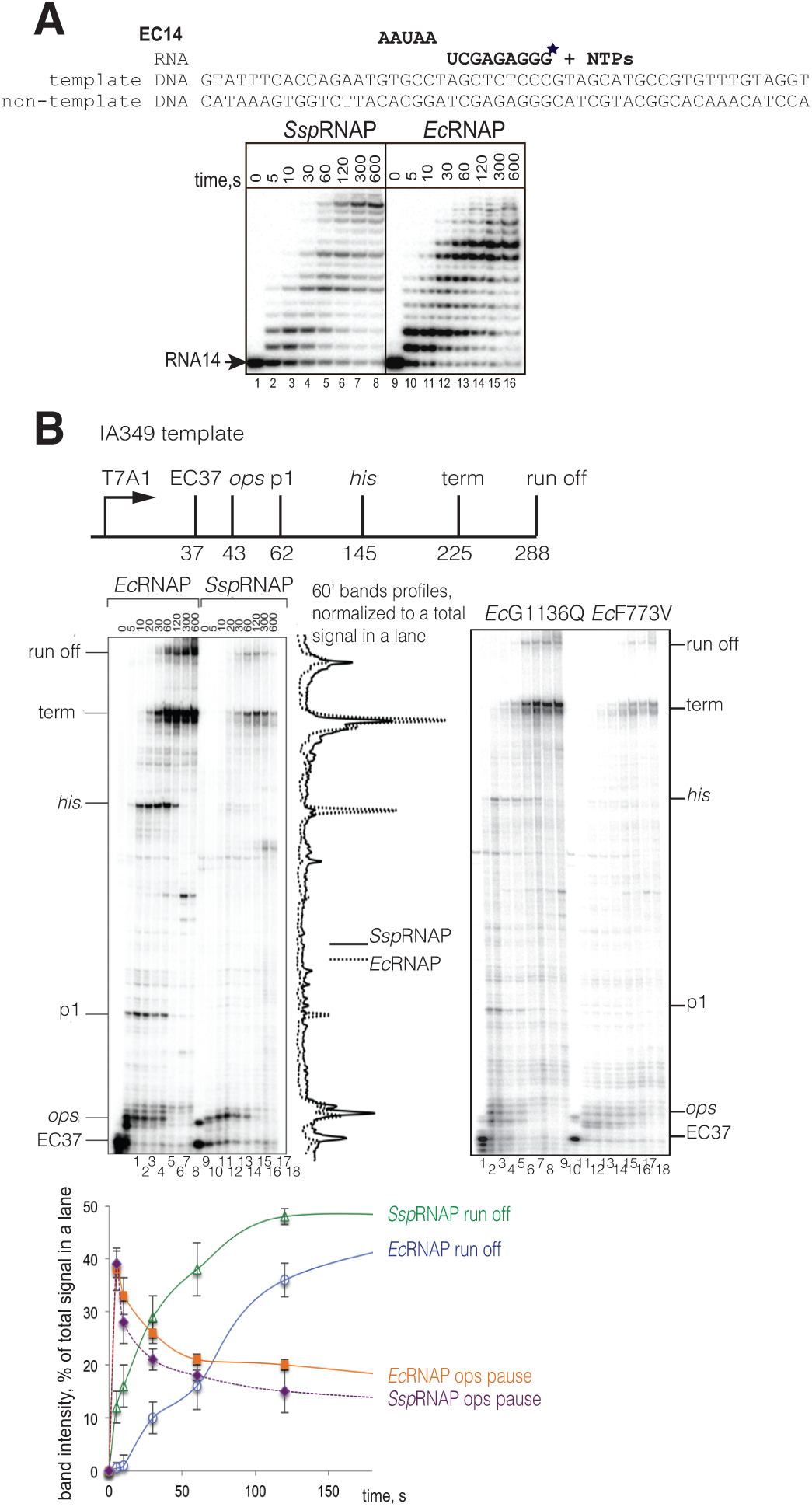
SspRNAP is less prone to pausing than EcRNAP. A. Kinetics of elongation in the EC14. (assembled elongation complex, scheme and sequence above the gel, asterisk indicates the position of ^32^P label) by SspRNAP and EcRNAPs with all four NTPs. **B. Single round elongation on IA349 template containing known pause sites and terminator**, shown on the scheme of the template above the gels images. Left gel - initial stalled elongation complex EC37 was formed with EcRNAP and SspRNAP, then chased with all four NTPs. To the right of the gel image, superimposed traces of the 60s bands for two RNAPs were generated by ImageQuant software and normalised to the total amount of radioactivity in the lane. To the left of the gel image a graph shows kinetics of run-off product accumulation and amount of complexes paused at ops pause sequence for EcRNAp and SspRNAP as a percentages from total radioactivity in the lane. Right gel shows kinetics of single round elongation on IA349 template performed similarly with mutant EcRNAPs with substitutions G1136Q and F773V.

To test if recognition of regulatory pausing and termination signals is similarly affected, we used the well-characterised template IA349, which contains an *ops, his*, an additional pause p1, and a rho-independent terminator sequence downstream of a T7A1 promoter (scheme in Fig. 6B). On a linear PCR-generated DNA template, after making a stalled elongation complexes with RNA37 in a subset of NTP substrates (Fig. 6B, left gel image, lanes 1 and 10), the kinetics of elongation at low concentration of NTPs were analysed. *Ssp*RNAP has an altered pausing pattern in comparison to *Ec*RNAP (Fig. 6B, compare the normalised traces from the 60 second time points). *Ssp*RNAP recognises the *ops* pause sequence, but not the *his* or p1 pauses; the efficiency of termination is also decreased. Similar behaviour is displayed by β’F773V and ‘G1136Q *E. coli* mutant RNAPs (Fig. 6B, right gel image), implying that amino acids in these positions possibly determine the *Ssp*RNAP’s pausing phenotype in general. Consistent with our findings, β’F773V was previously characterised as pause-resistant, and a different mutant in the 1136 position, β’G1136M, as fast elongating (33, 34, 37). The only pause recognised relatively efficiently by *Ssp*RNAP is the *ops* pause (plot with kinetics of *ops* pausing below gels), previously characterised as pretranslocated (29, 38). This result ascertains once more that *Ssp*RNAP ‘s equilibrium between pre- and post-translocation states is similar to that of *Ec*RNAP.

This experiment further confirmed that *Ssp*RNAP is more prone to reside in closed conformation, since neither the *his* pause nor terminator, which both require an open conformation of RNAP (31), are efficiently recognised by *Ssp*RNAP. Altogether our results suggest that *Ssp*RNAP is less responsive to diverse pausing signals, and transcription elongation in cyanobacteria might be more continuous process in general (see also Discussion) in comparison to that of *E. coli*. This property might further alleviate the need for proofreading factors in cyanobacteria.

## Discussion

Here, we found that cyanobacterial RNA polymerase efficiently performs the functions which are carried out by Gre/TFIIS factors in other taxons. Cyanobacterial RNAP possesses a very efficient intrinsic proofreading mechanism. This proofreading mechanism is potent enough to keep the rate of *in vivo* transcriptional msitakes in *Synechocystis sp* PCC 6803 at a level only fractionally lower than that of *E.coli*, and nevertheless low enough to be apparently easily tolerated.

Gre factors act at a molecular level by increasing affinity for the catalytic metal ion (18) and activating the attacking water (20). Affinity of the cyanobacterial enzyme for magnesium is not increased, consistent with conservation of amino acid residues in the close vicinity of catalytic magnesium. However, water deprotonation by *Ssp*RNAP is assisted by a general base in a similar fashion to GreA (20) (and *Thermus aquaticus* RNAP (15)). In contrast, in *E. coli*, a general base is not involved in hydrolysis, in agreement with the earlier suggestion of Mishanina et al. (17).

We propose that the cyanobacterial RNAP active site resides in a closed state by default, which results in suppression of pausing and termination. This conformation is thermodynamically related to a hydrolytically proficient conformation, reflected in a lower activation energy of the hydrolytic reaction. The closed conformation is fixed by amino acid residues in positions affecting the dynamics of the flexible TL and BH domains. Four amino acid residues specific to *Synechocystis sp* 6803 when introduced into *Ec*RNAP, significantly (3-11 times) increased the rate of hydrolysis, with magnitude A940V> F773V> G1136Q> N792D. Cyanobacteria and chloroplasts have non - *E. coli* amino acid residues at corresponding positions (Fig. 4A). The A940V change, never reported before, resulted in the highest acceleration of *in vitro* hydrolysis and the best complementation of a temperature sensitive phenotype of the double *greAgreB* deletion *E.coli* MG1655 strain. The second strongest effect has the F773V substitution, previously known to affect viability of *E.coli*, but never reported to affect hydrolysis. At position 1136 different changes (to S and M) were shown to increase rate of hydrolysis by *E. coli* and *D. radiodurans* RNAPs (26, 37), consistent with our results.

These residues are located too far from the active site to directly participate in the reaction, but they might change the conformation and flexibility of the TL and BH. G1136 is located in the hinge region of the TL, A940V is next to a double glycine motif at the tip of the TL (Fig. 4B, flexible parts are highlighted in cyan). In support of our hypothesis, substitutions F773V and G1136S in *E. coli* were predicted to stabilise a closed TL (39, 40) and recently were shown to stabilise a 1bp backtracked state (41). Similarly, residue N792 is located in the flexible hinge region of the BH, and mutants at this position were predicted to fix the BH in a particular conformation (42, 43). We cannot completely rule out input from other domains of RNAP into efficient proofreading; additional specific features may contribute to cyanobacterial RNAP protein dynamics, such as the physical location of the catalytic aspartate triad and TL/BH modules on two separate proteins due to the split of β’subunit and the presence of a much larger SI3 insertion.

Importantly, although isomerisation of *Ssp*RNAP into a 1bp backtracked state is very efficient, longer backtracking has a similar rate to *Ec*RNAP, meaning that there is no general propensity of cyanobacterial RNAP to move backwards and, therefore, no risk of frequent elongation interruptions. Perhaps, there are two separate thermodynamic or physical routes for 1bp backtracking vs longer backtracking, and only one is changed in cyanobacteria.

Another consequence of the cyanobacterial active site conformation is a reduced ability of RNAP to regulate elongation via pausing. This loss is a big sacrifice, since RNAP pausing in bacteria and eukaryotes is a major regulatory mechanism, coupling transcription with other cellular processes (44). Cyanobacteria might either lack this regulatory possibility entirely or employ as yet unknown regulatory factors. The high efficiency of hydrolysis and fast elongation are common for two distantly related species of cyanobacteria, *Synechocystis sp* PCC 6803 and *Synechococcus elongatus* 7942, suggesting that these are general features of cyanobacterial transcription. Plant chloroplasts (descendants of ancient cyanobacteria) and other groups that lack proofreading factors, might share these features.

What are the physiological benefits of cyanobacterial reliance on intrinsic proofreading? It solves the problem of recruitment of proofreading factors, and the cyanobacterial set up is reminiscent of eukaryotic RNAPs I and III, which carry out proofreading activity on one of the subunits (45). Notably, the hydrolytic activity of *Ssp*RNAP decreased with decreasing temperature at a slower rate than in *E. coli*, and was still apparent even at zero degrees, suggesting that in low winter temperatures intrinsic proofreading is still active in cyanobacteria when protein diffusion is slow (Fig. S4).

We assume that Gre factors were never acquired by cyanobacteria in evolution. Transcript cleavage factors are not conserved between the Archaea /Eukarya and Bacteria (46). According to the recently published tree of life, cyanobacteria belong to a deep phylum stemming directly from a common ancestor of Archaea /Eukarya and Bacteria; meaning that they might have branched out before proofreading factors acquisition (47).

Alternatively, Gre factors might have been lost after they became dispensable. The ultimate cellular role of Gre factors is to alleviate conflicts between the replication and transcription machineries (25), which may occur less frequently in cyanobacteria due to circadian regulation and polyploidy. In photosynthetic cyanobacteria, replication is thought not to coincide in time with the main peak of transcription (48). RNAP and the replisome might not operate on the same copy of genome in polyploidal cyanobacteria (e.g. *Synechocystis sp* 6803 has more than 10 copies of its genome in a cell (49)). Hypothetically, Gre factors could have become detrimental because they can potentially bring a wrong metal ion, such as iron (very abundant in cyanobacteria (50)), into RNAP active site, which when combined with reactive oxygen species generated in by the electron transport chain, leads to protein damage (51). Gre factors interference with both nucleotide excision (52) and double strand break repair (53), are another possible incentives for their loss from cyanobacteria, which have a high level of genomic DNA photodamage. Acquisition of an extremely large SI3 (∼650 aa) could also led to occlusion of the Gre binding site and sped up the loss.

## Material and Methods

### Strains and plasmids

*Synechocystis sp* PCC 6803 strain was a gift from Prof Robinson, Durham University, UK; *Synechococcus elongatus* PCC 7942 was a gift from Prof Mullineaux, Queen Mary University of London; *E. coli* strain PGe74 (MG1655 *ΔgreAgreB*) was a gift from Dr P.Gamba, Newcastle University, UK. Plasmid pIA349 was a gift from Prof. I.Artsimovitch, Ohio State University, USA.

### Mutagenesis and RNAPs purification

To generate mutation in *E.coli* rpoC gene, pRL663 plasmid encoding *E. coli* rpoC with C-terminal His_6_-tag under IPTG inducible promoter was used as template for mutagenesis by QuickChange XL Mutagenesis Kit, ThermoFisher, according to the manufacturers’ protocol. WT and mutant constructs were transformed into *E. coli* MG1655 *ΔgreAgreB* strain. Cell cultures were grown up to OD_600_ of 0.6, and 1mM IPTG was added to induce expression of plasmid-borne rpoC for 3.5 hours. RNAPs were purified by Heparin (HiTrap Heparin column, GE Healthcare), Ni-NTA affinity (on HisTrap column, GE Healthcare) and ion exchange (ResourseQ column, GE Healthcare) chromatographic steps. *Synechocystis sp* PCC 6803 was grown at constant light (100 µmol photons m^−2^ s^−1^) at 30°C in BG-11. Harvested cells were disrupted using bead beater with 0.1mm zirconia beads. After centrifugation at 15000 rpm for 20 min, followed by ultracentrifugation at 100000 rpm for 1hr (to remove membrane fraction), lysate was loaded onto Heparin column (HiTrap Heparin column, GE Healthcare). RNAP was further purified using size exclusion (Superose 6, GE Healthcare) and ion-exchange chromatography (ResourseQ column, GE Healthcare) steps.

### Transcription Assays

All transcription experiments were done at 30°C in transcription buffer containing 20 mM Tris-HCl pH 7, 40 mM KCl, 10 mM MgCl_2_, unless otherwise specified. Elongation complexes were assembled and immobilised on streptavidin sepharose beads (GE Healthcare) as described (14, 16). Sequences of the oligonucleotides used for the elongation complexes assembly are shown either in corresponding Figures or Fig. S1. RNA was either kinased at the 5’-end using [γ-^32^P] ATP or labelled at the 3’-end after elongation complex assembly by incorporation of [α-^32^P] GTP (Hartmann Analytic), dictated by template, with subsequent removal of unincorporated nucleotide by washing beads with transcription buffer. To determine the rate of nucleotide addition, 1 μM NTP together with 10 mM MgCl_2_ (final concentrations), was added to initial EC, reactions were incubated at room temperature and stopped by addition of formamide /8M urea - containing loading buffer. Products were resolved by denaturing (8 M urea) 23% PAGE, revealed by PhosphorImaging (GE Healthcare) and analysed using ImageQuant software (GE Healthcare). The proportion of elongated RNA was plotted against time and fitted to a single exponential equation by using nonlinear regression in SigmaPlot software.

Misincorporation was initiated by addition of a mixture 10 mM MgCl_2_, 1mM non-cognate NTP, reactions were kept at 30°C for times indicated in the Figure 2A. To determine the rate of misincorporation, the proportion of complexes that undergone misincorporation (and subsequent cleavage) was plotted against time and fitted as above. Cleavage reactions were initiated by addition of 10 mM MgCl_2_ (final concentration), unless otherwise specified. Reactions were incubated at 30°C for times indicated in the figures, and were stopped by addition of formamide/8M urea - containing loading buffer. Products were resolved by denaturing (8 M urea) 23% PAGE, revealed by PhosphorImaging (GE Healthcare) and analysed using ImageQuant software (GE Healthcare). To determine the rate of phosphodiester bond hydrolysis proportion of the cleaved RNA was plotted against time and fitted to a single exponential equation using non-linear regression (14, 16). To determine the k_M_ (Mg^2+^) for cleavage in mEC(A) and mEC(U) the reaction rates obtained at various MgCl_2_ concentrations were fitted to the Michaelis-Menten equation (14, 16). For activation energy calculation, cleavage rates in mEC(U) at 0, 5,10, 15, 20, 25, 30, 37, 42 and 45°C were calculated first and then plotted in lnK vs 1/T coordinates. Data were fitted into linear equation lnK=lnA-E_a_/R(1/T) using SigmaPlot software.

For elongation experiments on IA349 template initial stalled complex EC37 was formed using 150 μM CAUC, 5 μM ATP, 5 μM CTP, 1.3 μM [α-^32^P] GTP (700 Ci/mmol) and biotinylated template ds PCR generated template DNA. After 2 minutes incubation, streptavidine sepharose (GE Healthcare) beads were added and incubated for additional 2 minutes, then washed twice with transcription buffer with 200 mM NaCl and twice with transcription buffer. Elongation was started by addition of 1g mM NTPs and 10 mM MgCl_2_.

### Growth complementation assay

*E. coli* strain MG1655 *ΔgreAgreB* was transformed with pRL663 (54) based plasmids expressing either WT or mutant β’ subunits under IPTG inducible bacterial promoter. Overnight cultures were diluted and grown until mid-exponential phase, then each culture was diluted up to OD_600_ 0.1. These initial cultures were further serially diluted, plated and incubated overnight either at 30°C or 37°C with addition of 0.1 mM IPTG to drive the expression of plasmid-encoded subunit.

### Next Generation Sequencing and data analysis

Total RNA was isolated from mid-exponentially growing cultures of *E. coli* and *Synechocystis sp* PCC 6803 as described in (55). Quality of RNA was checked by BioAnalyser, sample preparation and sequencing were performed by Vertis Ltd, essentially as described in (24), the only modification of the protocol is usage of PrimeScript, Clontech high fidelity reverse transcriptase. Dataset quality was assessed using FastQC (https://www.bioinformatics.babraham.ac.uk/projects/fastqc/) to ensure per base and per tile sequence quality. Raw reads were trimmed using fastx_trimmer (http://hannonlab.cshl.edu/fastx_toolkit/). Trimmed reads were aligned to genomes using Bowtie (56) allowing three mismatched bases and only unique alignments (-n 3 -m 1). *E. coli* alignment used the NC_000913.3 reference genome and *Synechocystis* 6803 used the consensus derived from the in house sequencing data using CLC Workbench (https://www.qiagenbioinformatics.com/). Single base variations between the experimental *E. coli* strain and the reference genome were identified using samtools and bcftools (57). Error rate analysis was carried out in R using the BioConductor seqTools, seqInR and IRanges packages (58, 59). Total error rates were calculated as the percentage of total reads with a mismatched base at each read position in the alignment, thresholded to a Phred quality score of <30. Specific error rates were calculated as the percentage of total reads with a specific mismatch, for example an A incorporated instead of a G (G > A misincorporation), at each read position 7, thresholded to a Phred quality score of <30. Ambiguous N bases and positions of single base variation were excluded from the error rate calculation. Raw and processed data were uploaded into GEO Database, under accession number GSE115135.

## Data availability

Data generated from analysis of NGS in both raw and processed form are available at GEO Database, under accession number GSE115135.

## Acknowledgements

The work was supported by the UK Biotechnology and Biological Sciences Research Council DTP Studentship (A.R.B.), Royal Society Fellowship and EPSRC grant EP/N031962/1 (Y.Y.)

## Author contributions

A.R.B. conceived a number of experiments, conducted and analysed the results of most of the experiments, K.J. analysed NGS data, Y.Y. conceived most of the experiments and conducted some, analysed data and wrote a manuscript.

## References

1. Libby RT & Gallant JA (1991) The role of RNA polymerase in transcriptional fidelity. Mol Microbiol 5(5):999–1004.

2. Nielsen S & Zenkin N (2013) Transcript assisted phosphodiester bond hydrolysis by eukaryotic RNA polymerase II. Transcription 4(5).

3. Orlova M, Newlands J, Das A, Goldfarb A, & Borukhov S (1995) Intrinsic transcript cleavage activity of RNA polymerase. Proc Natl Acad Sci U S A 92(10):4596–4600.

4. Sigurdsson S, Dirac-Svejstrup AB, & Svejstrup JQ (2010) Evidence that transcript cleavage is essential for RNA polymerase II transcription and cell viability. Mol Cell 38(2):202–210.

5. Borukhov S, Polyakov A, Nikiforov V, & Goldfarb A (1992) GreA protein: a transcription elongation factor from Escherichia coli. Proc Natl Acad Sci U S A 89(19):8899–8902.

6. Borukhov S, Sagitov V, Josaitis CA, Gourse RL, & Goldfarb A (1993) Two modes of transcription initiation in vitro at the rrnB P1 promoter of Escherichia coli. J. Biol. Chem 268:23477–23482.

7. Jeon C & Agarwal K (1996) Fidelity of RNA polymerase II transcription controlled by elongation factor TFIIS. Proc Natl Acad Sci U S A 93(24):13677–13682.

8. Archambault J, Lacroute F, Ruet A, & Friesen JD (1992) Genetic interaction between transcription elongation factor TFIIS and RNA polymerase II. Mol Cell Biol 12(9):4142–4152.

9. Lange U & Hausner W (2004) Transcriptional fidelity and proofreading in Archaea and implications for the mechanism of TFS-induced RNA cleavage. Mol Microbiol 52(4):1133–1143.

10. Steitz TA (1998) A mechanism for all polymerases. Nature 391(6664):231–232.

11. Wang B, Predeus AV, Burton ZF, & Feig M (2013) Energetic and structural details of the trigger-loop closing transition in RNA polymerase II. Biophys J 105(3):767–775.

12. Zhang J, Palangat M, & Landick R (2009) Role of the RNA polymerase trigger loop in catalysis and pausing. Nat Struct Mol Biol 17(1):99–104.

13. Toulokhonov I, Zhang J, Palangat M, & Landick R (2007) A central role of the RNA polymerase trigger loop in active-site rearrangement during transcriptional pausing. Mol Cell 27(3):406–419.

14. Yuzenkova Y, et al. (2010) Stepwise mechanism for transcription fidelity. BMC Biol 8:54.

15. Yuzenkova Y & Zenkin N (2010) Central role of the RNA polymerase trigger loop in intrinsic RNA hydrolysis. Proc Natl Acad Sci U S A 107(24):10878–10883.

16. Zenkin N, Yuzenkova Y, & Severinov K (2006) Transcript-assisted transcriptional proofreading. Science 313(5786):518–520.

17. Mishanina TV, Palo MZ, Nayak D, Mooney RA, & Landick R (2017) Trigger loop of RNA polymerase is a positional, not acid-base, catalyst for both transcription and proofreading. Proc Natl Acad Sci U S A 114(26):E5103–E5112.

18. Laptenko O, Lee J, Lomakin I, & Borukhov S (2003) Transcript cleavage factors GreA and GreB act as transient catalytic components of RNA polymerase. Embo J 22(23):6322–6334.

19. Roghanian M, Yuzenkova Y, & Zenkin N (2011) Controlled interplay between trigger loop and Gre factor in the RNA polymerase active centre. Nucleic Acids Res 39(10):4352–4359.

20. Sosunova E, Sosunov V, Epshtein V, Nikiforov V, & Mustaev A (2013) Control of transcriptional fidelity by active center tuning as derived from RNA polymerase endonuclease reaction. J Biol Chem 288(9):6688–6703.

21. Bubunenko MG, et al. (2017) A Cre Transcription Fidelity Reporter Identifies GreA as a Major RNA Proofreading Factor in Escherichia coli. Genetics 206(1):179–187.

22. Marr MT & Roberts JW (2000) Function of transcription cleavage factors GreA and GreB at a regulatory pause site. Mol Cell 6(6):1275–1285.

23. Borukhov S, Sagitov V, & Goldfarb A (1993) Transcript cleavage factors from E. coli. Cell 72(3):459–466.

24. Yuzenkova Y, et al. (2014) Control of transcription elongation by GreA determines rate of gene expression in Streptococcus pneumoniae. Nucleic Acids Res 42(17):10987–10999.

25. Trautinger BW, Jaktaji RP, Rusakova E, & Lloyd RG (2005) RNA polymerase modulators and DNA repair activities resolve conflicts between DNA replication and transcription. Mol Cell 19(2):247–258.

26. Esyunina D, et al. (2016) Lineage-specific variations in the trigger loop modulate RNA proofreading by bacterial RNA polymerases. Nucleic Acids Res 44(3):1298–1308.

27. Imashimizu M, Oshima T, Lubkowska L, & Kashlev M (2013) Direct assessment of transcription fidelity by high-resolution RNA sequencing. Nucleic Acids Res.

28. Imashimizu M, Tanaka K, & Shimamoto N (2011) Comparative Study of Cyanobacterial and E. coli RNA Polymerases: Misincorporation, Abortive Transcription, and Dependence on Divalent Cations. Genet Res Int 2011:572689.

29. Artsimovitch I & Landick R (2000) Pausing by bacterial RNA polymerase is mediated by mechanistically distinct classes of signals. Proc Natl Acad Sci U S A 97(13):7090–7095.

30. Komissarova N & Kashlev M (1997) Transcriptional arrest: Escherichia coli RNA polymerase translocates backward, leaving the 3’ end of the RNA intact and extruded. Proc Natl Acad Sci U S A 94(5):1755–1760.

31. Sekine S, Murayama Y, Svetlov V, Nudler E, & Yokoyama S (2015) The ratcheted and ratchetable structural states of RNA polymerase underlie multiple transcriptional functions. Mol Cell 57(3):408–421.

32. Zhang N, et al. (2015) Mutations in RNA Polymerase Bridge Helix and Switch Regions Affect Active-Site Networks and Transcript-Assisted Hydrolysis. J Mol Biol 427(22):3516–3526.

33. Artsimovitch I, Chu C, Lynch AS, & Landick R (2003) A new class of bacterial RNA polymerase inhibitor affects nucleotide addition. Science 302(5645):650–654.

34. Nedialkov YA, et al. (2013) The RNA polymerase bridge helix YFI motif in catalysis, fidelity and translocation. Biochim Biophys Acta 1829(2):187–198.

35. Larson MH, et al. (2012) Trigger loop dynamics mediate the balance between the transcriptional fidelity and speed of RNA polymerase II. Proc Natl Acad Sci U S A 109(17):6555–6560.

36. Weixlbaumer A, Leon K, Landick R, & Darst SA (2013) Structural basis of transcriptional pausing in bacteria. Cell 152(3):431–441.

37. Bar-Nahum G, et al. (2005) A ratchet mechanism of transcription elongation and its control. Cell 120(2):183–193.

38. Bochkareva A, Yuzenkova Y, Tadigotla VR, & Zenkin N (2012) Factor-independent transcription pausing caused by recognition of the RNA-DNA hybrid sequence. Embo J 31(3):630–639.

39. Mejia YX, Nudler E, & Bustamante C (2015) Trigger loop folding determines transcription rate of Escherichia coli’s RNA polymerase. Proc Natl Acad Sci U S A 112(3):743–748.

40. Malinen AM, et al. (2014) CBR antimicrobials alter coupling between the bridge helix and the beta subunit in RNA polymerase. Nat Commun 5:3408.

41. Turtola M, Makinen JJ, & Belogurov GA (2018) Active site closure stabilizes the backtracked state of RNA polymerase. Nucleic Acids Res 46(20):10870–10887.

42. Tuske S, et al. (2005) Inhibition of bacterial RNA polymerase by streptolydigin: stabilization of a straight-bridge-helix active-center conformation. Cell 122(4):541–552.

43. Tan L, Wiesler S, Trzaska D, Carney HC, & Weinzierl RO (2008) Bridge helix and trigger loop perturbations generate superactive RNA polymerases. J Biol 7(10):40.

44. Landick R (2006) The regulatory roles and mechanism of transcriptional pausing. Biochem Soc Trans 34(Pt 6):1062–1066.

45. Alic N, et al. (2007) Selectivity and proofreading both contribute significantly to the fidelity of RNA polymerase III transcription. Proc Natl Acad Sci U S A 104(25):10400–10405.

46. Werner F & Grohmann D (2011) Evolution of multisubunit RNA polymerases in the three domains of life. Nat Rev Microbiol 9(2):85–98.

47. Hug LA, et al. (2016) A new view of the tree of life. Nat Microbiol 1:16048.

48. Paijmans J, Bosman M, Ten Wolde PR, & Lubensky DK (2016) Discrete gene replication events drive coupling between the cell cycle and circadian clocks. Proc Natl Acad Sci U S A 113(15):4063–4068.

49. Griese M, Lange C, & Soppa J (2011) Ploidy in cyanobacteria. FEMS Microbiol Lett 323(2):124–131.

50. Shcolnick S & Keren N (2006) Metal homeostasis in cyanobacteria and chloroplasts. Balancing benefits and risks to the photosynthetic apparatus. Plant Physiol 141(3):805–810.

51. Patterson CO & Myers J (1973) Photosynthetic Production of Hydrogen Peroxide by Anacystis nidulans. Plant Physiol 51(1):104–109.

52. Epshtein V, et al. (2014) UvrD facilitates DNA repair by pulling RNA polymerase backwards. Nature 505(7483):372–377.

53. Sivaramakrishnan P, et al. (2017) The transcription fidelity factor GreA impedes DNA break repair. Nature 550(7675):214–218.

54. Yuzenkova J, et al. (2002) Mutations of bacterial RNA polymerase leading to resistance to microcin j25. J Biol Chem 277(52):50867–50875.

55. Forrest D, James K, Yuzenkova Y, & Zenkin N (2017) Single-peptide DNA-dependent RNA polymerase homologous to multi-subunit RNA polymerase. Nat Commun 8:15774.

56. Langmead B, Trapnell C, Pop M, & Salzberg SL (2009) Ultrafast and memory-efficient alignment of short DNA sequences to the human genome. Genome biology 10(3):R25.

57. Li H, et al. (2009) The Sequence Alignment/Map format and SAMtools. Bioinformatics 25(16):2078–2079.

58. Ugo Bastolla MP, H. Eduardo Roman, Michele Vendruscolo (2007) Structural Approaches to Sequence Evolution (Springer Verlag).

59. Lawrence M, et al. (2013) Software for computing and annotating genomic ranges. PLoS computational biology 9(8):e1003118.

