## Supplemental text and figures for "RNA polymerase active centre compensates for the absence of transcription proofreading factors in Cyanobacteria"

**template 2**

**cEC(A)**

```
                              AAUAA
          RNA15(A)                   UCGAGAGGGA*
      template DNA        TGAATGTAGGTAGCTCTCCCTGTGCCGCTTATCGGT
  non-template DNA        ACTTACAGCCATCGAGAGGGACACGGCGAATAGCCA
```

**mEC(A)**

```
                             *AAUAA                    A
          RNA15(A)                   UCGAGAGGG
      template DNA        TGAATGTAGGTAGCTCTCCCCGTGCCGCTTATCGGT
  non-template DNA        ACTTACAGCCATCGAGAGGGGCACGGCGAATAGCCA
```

**mEC(U)**

```
                             *AAUAA                    U
          RNA15(U)                   UCGAGAGGG
      template DNA        TGAATGTAGGTAGCTCTCCCGGTGCCGCTTATCGGT
  non-template DNA        ACTTACAGCCATCGAGAGGGCCACGGCGAATAGCCA
```

**mEC(C)**

```
                             *AAUAA                    C
          RNA15(C)                   UCGAGAGGG
      template DNA        TGAATGTAGGTAGCTCTCCCAGTGCCGCTTATCGGT
  non-template DNA        ACTTACAGCCATCGAGAGGGTCACGGCGAATAGCCA
```

**Fig. S.1. Structure of elongation complexes based on template 2, and sequences of nucleic acid oligonucleotides used for their assembly.** Asterisk indicates that RNA is labelled at the 5' - end by kinasing oligonucleotide using polynucleotide kinase A and [γ- $P^{32}$] ATP.

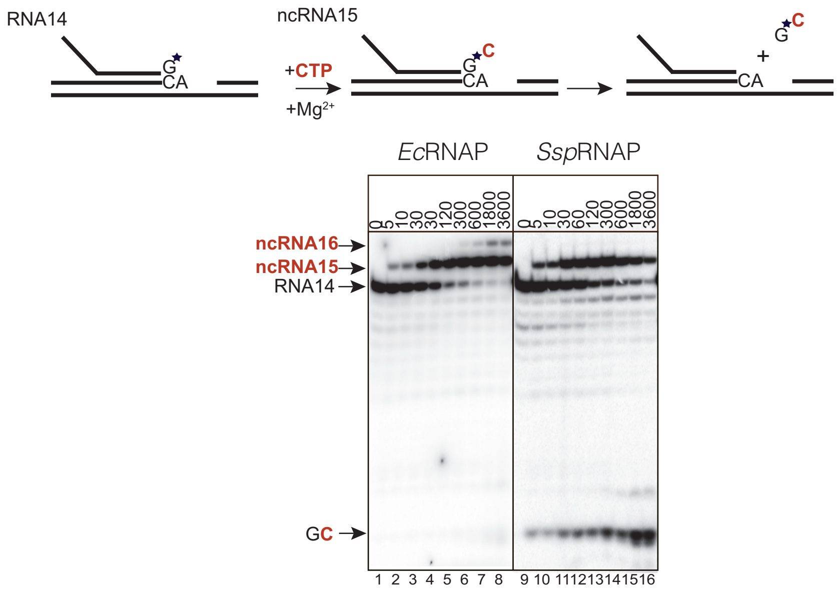


**Fig. S. 2. Kinetics of misincorporation of CTP instead of UTP** dictated by template with concomitant proofreading reaction - cleavage of the GC 3' dinucleotide portion of the transcript by EcRNAp and SspRNAP. Above the gel is the scheme of reaction; asterisk indicates that RNA is labelled at the 3' end by incorporation of radiolabelled GTP into the transcript. *Ssp*RNAP).

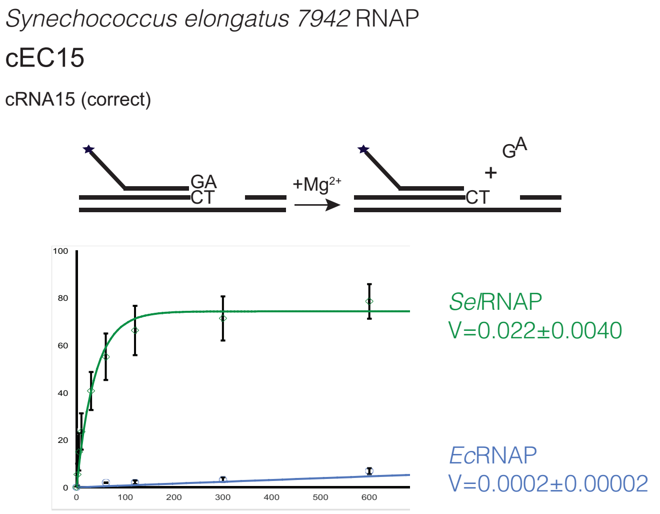


**Fig. S.3. Kinetics of transcript hydrolysis in the correct elongation complex cEC(A) (Fig. S1) by purified RNAP of different cyanobacterium, Synechococcus elongatus, *Sel*RNAP.** Above the gel is the scheme of reaction; asterisk indicates that RNA is labelled at the 5’ - end by kinasing 13 nt RNA oligo using polynucleotide kinase A and [γ- $P^{32}$] ATP. After assembly into elongation complex, initial RNA13 was extended to RNA15 with addition of ATP and GTP. The graph fits of data to the exponential equation are shown as solid lines, *Ec*RNAP (blue) and *Sel*RNAP (green).

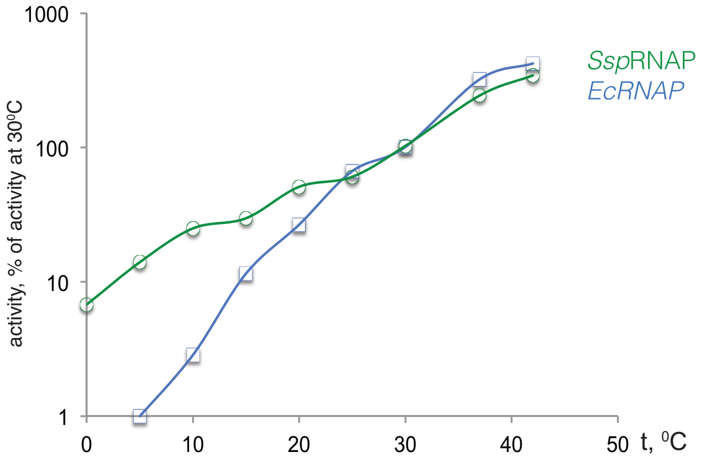


**Fig. S.4. Temperature –dependence of rate of hydrolysis reaction in mEC(U) by *Ec*RNAP and *Ssp*RNAP.**

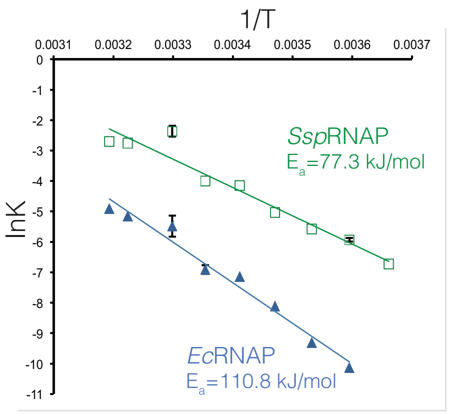


**Fig. S.5. Activation energy for RNA hydrolysis in mEC(A) elongation complex by *Ec*RNAP and *Ssp*RNAP.** . Arrhenius plots for *Ec*RNAP and *Ssp*RNAP, graph fit of lnK to 1/T data to linear equation are shown as straight line, apparent activation energy calculated from equation $\ln K = \ln A - E_a/R(1/T)$ is shown on the plot. The data points are averages of two independent experiments.

**Table S1. Rates of transcript hydrolysis in mEC(A) and mEC(U) by WT and mutant *Ec*RNAPs with indicated amino acid changes in β' subunit at 30$^0$C and 10 $MgCl_2$.**

| RNAP | mEC(U) hydrolysis rate, s-1 | Standart deviation | mEC(A) hydrolysis rate, s-1 | Standart deviation | mEC(C) hydrolysis rate, s-1 | Standart deviation | Ratio mutant/wt mEC(U) | Ratio mutant/wt mEC(A) |
|---|---|---|---|---|---|---|---|---|
| *Ec* WT | 0.0010 | 0.00021 | 0.0035 | 0.00064 | 0.0005 | 0.00006 | 1 | 1 |
| *Ec* Q771E | 0.0014 | 0.00107 | 0.0042 | 0.00074 | | | 1.5 | 1.2 |
| *Ec* F773V | 0.0082 | 0.00165 | 0.0290 | 0.00232 | 0.0043 | 0.00030 | 8.6 | 8.1 |
| *Ec* N792D | 0.0063 | 0.00056 | 0.0125 | 0.00195 | | | 6.6 | 3.5 |
| *Ec* I937T | 0.0007 | 0.00031 | 0.0039 | 0.00071 | | | 0.8 | 1.1 |
| *Ec* A940V | 0.0105 | 0.0010 | 0.0260 | 0.00532 | | | 11.1 | 7.3 |
| *Ec* A941F | 0.0003 | 0.00004 | 0.0011 | 0.00013 | | | 0.3 | 0.3 |
| *Ec* K1134G | 0.0011 | 0.0002 | 0.0029 | 0.0004 | | | 1.2 | 0.8 |
| *Ec* G1136Q | 0.0056 | 0.00103 | 0.0111 | 0.00272 | 0.0023 | 0.00030 | 5.8 | 3.1 |
| *Ec* V1141I | 0.0001 | 0.00002 | 0.0056 | 0.00072 | | | 1.1 | 1.5 |
| *Ec* A1142E | 0.0007 | 0.0001 | 0.0082 | 0.00122 | | | 0.7 | 2.3 |
| *Ec* D1143E | 0.0012 | 0.0002 | 0.0051 | 0.00073 | | | 1.2 | 1.4 |
| *Ec* F1145L | 0.0028 | 0.00074 | 0.0033 | 0.00121 | | | 2.9 | 0.9 |
| SspRNAP | 0.0612 | 0.00814 | 0.1401 | 0.02423 | | | 63.1 | 39.3 |
| SelRNAP | 0.0502 | 0.00782 | 0.1212 | 0.00735 | | | 52.6 | 33.7 |
